# Antibiotic resistant and extended-spectrum β-lactamase producing faecal coliforms in wastewater treatment plant effluent

**DOI:** 10.1101/839399

**Authors:** Cian Smyth, Aidan O’Flaherty, Fiona Walsh, Thi Thuy Do

**Affiliations:** Department of Biology, Maynooth University, Maynooth, Co. Kildare, Ireland

**Keywords:** Antibiotic resistance, multidrug resistance, AmpC, extended-spectrum β-lactamase (ESBL), plasmids

## Abstract

Wastewater treatment plants (WWTPs) provide optimal conditions for the maintenance and spread of antibiotic resistant bacteria (ARB) and antibiotic resistance genes (ARGs). In this work we describe the occurrence of antibiotic resistant faecal coliforms and their mechanisms of antibiotic resistance in the effluent of two urban WWTPs in Ireland. Effluent samples were collected from two WWTPs in Spring and Autumn of 2015 and 2016. The bacterial susceptibility patterns to 13 antibiotics were determined. The phenotypic tests were carried out to identify AmpC or extended-spectrum β-lactamase (ESBL) producers. The presence of ESBL genes were detected by PCR. Plasmids carrying ESBL genes were transformed into *Escherichia coli* DH5α recipient and underwent plasmid replicon typing to identify incompatibility groups. More than 90% of isolated faecal coliforms were resistant to amoxicillin and ampicillin, followed by tetracycline (up to 39.82%), ciprofloxacin (up to 31.42%) and trimethoprim (up to 37.61%). Faecal coliforms resistant to colistin and imipenem were detected in all effluent samples. Up to 53.98% of isolated faecal coliforms expressed a multi-drug resistance (MRD) phenotype. AmpC production was confirmed in 5.22% of isolates. The ESBL genes were confirmed for 11.84% of isolates (9.2% of isolates carried *bla*_TEM_, 1.4% *bla*_SHV-12_, 0.2% *bla*_CTX-M-1_ and 1% *bla*_CTX-M-15_). Plasmids extracted from 52 ESBL isolates were successfully transformed into recipient *E. coli*. The detected plasmid incompatibility groups included the IncF group, IncI1, IncHI1/2 and IncA/C. These results provide evidence that treated wastewater is polluted with ARB and MDR faecal coliforms and are sources of ESBL-producing, carbapenem and colistin resistant *Enterobacteriaceae*.

**Importance:** Antibiotic resistant bacteria (ARB) are an emerging environmental concern with a potential impact on human health. The results provide the evidence that treated wastewater is polluted with antibiotic resistant bacteria containing mobile resistance mechanisms of importance to clinical treatment of pathogens and multi-drug resistant (MDR) faecal coliforms. They are sources of relatively high proportions of ESBL-producing *Enterobacteriaceae*, and include carbapenem and colistin resistant *Enterobacteriaceae.* The significance of this study is the identification of the role of WWTPs as a potential control point to reduce or stop the movement of ESBL, MDR and colistin resistant bacteria into the environment from further upstream sources, such as human or animal waste.

## Introduction

The dissemination of antimicrobial resistance within bacterial communities and the selection of new resistance mechanisms are due to the large-scale use of antibiotics in agricultural, veterinary and human clinical applications (1–8). The emergence of antibiotic resistant bacteria (ARB) is a major public health issue which poses a serious therapeutic challenge worldwide (6). Wastewater treatment plants (WWTPs) are considered potential sources of ARB and antibiotic resistance genes (ARGs), which play an important role in the spread of antibiotic resistance into the environment (9, 10). The urban WWTPs receive wastewater from human communities, which contains high concentrations of chemical matter, including antibiotics and microorganisms, including ARB. Therefore, WWTPs are favourable environments with optimal conditions for the development and spread of ARB and ARGs (11, 12). Both ARB and ARGs were detected in wastewater samples globally (13–24). However, little is known of the fate of these bacteria; and the role of WWTPs in releasing ABR and ARGs into the environment through treated effluent (13). A recent report by Flach et al shows no evidence for the selection of antibiotic resistance in WWTPs (25); however, large amounts of resistant bacteria were identified throughout the wastewater treatment process (7, 26). The conventional wastewater treatment process can remove some ARB (27, 28), but ARB were still found in large proportions in the effluent (18, 29). In some cases, ARB were detected at higher rates in WWTP effluent than in the influent (30, 31).

AmpC cephalosporinases and extended-spectrum β-lactamases (ESBLs) are some of the most clinically important antibiotic resistance mechanisms (32). The dissemination of AmpC or ESBL producing *Escherichia coli* were identified in different types of aquatic environments, particularly in wastewater (33–36). The prevalence of AmpC or ESBL producing bacteria pose a global health problem due to limitations of therapeutic options for the treatment of infections caused by these bacteria (37). The ESBL genes are frequently located on mobile genetic elements (38). Among more than 300 subtypes of ESBL genes, *bla*_TEM_ and *bla*_SHV_ groups were the most common ESBL genes identified in human pathogens until the late 1990s. These groups were replaced by *bla*_CTX-M_ genes since the beginning of the 2000s and *Escherichia coli* became the most prevalent ESBL producing bacteria among clinical *Enterobacteriaceae* (39).

The monitoring of antibiotic resistance from WWTPs provides the information required to track the dissemination of ARB and ARGs into the environment (40, 41). Moreover, the analysis of ARB and ARGs in urban WWTPs is considered as an alternative method for the indirect study of antibiotic resistance in human populations from which the WWTPs receive wastewater (42). Indeed, the resistance rates of indicator bacteria in wastewater may give useful information to identify the changes in resistance in the human populations (43). The objectives of this study were to characterise the faecal coliforms resistome leaving urban WWTPs via the effluent. This was achieved through i) assessment of the prevalence of antibiotic resistant faecal coliforms in the effluent from two Irish urban WWTPs, ii) characterisation of the antibiotic resistance profiles of these bacteria, iii) identification of the occurrence of AmpC or ESBL producing faecal coliforms and iv) identification of the resistance mechanisms and their potential mobility.

## Materials and Methods

### Isolation of total faecal coliforms

Final effluent samples were collected from two urban WWTPs in Ireland during early Spring and late Autumn in 2015 and 2016. These WWTPs were representative of medium sized WWTPs with 100% urban agglomerations, include tertiary treatment, and the distance between them was less than 100km. Faecal coliforms were isolated using the membrane filtration method (44). The effluent samples (1 mL and 10 mL) were filtered through nitrocellulose membranes (Sigma Aldrich/Merck). The filters were then incubated on mFC agar at 37 °C for 24 hours in the presence or absence of antibiotics: amoxicillin (32 mg/L), ciprofloxacin (1 mg/L) or tetracycline (16 mg/L). All procedures were performed in triplicate.

### Antibiotic susceptibility test using agar dilution and disk diffusion methods

Bacterial isolates were subjected to antibiotic susceptibility testing using the agar dilution method following the CLSI recommendations (45). The antibiotics tested were tetracycline, amoxicillin, ampicillin, ciprofloxacin, kanamycin, gentamicin, colistin, chloramphenicol and trimethoprim. The imipenem, meropenem, cefotaxime and ceftazidime susceptibilities were determined using the disk diffusion method (45). The minimum inhibitory concentration (MIC) breakpoints for Enterobacteriaceae in the CLSI guidelines (45) were used to identify ARB. The EUCAST MIC breakpoint for colistin was used (46). Bacterial isolates resistant to three or more different antibiotic classes of antibiotics were defined as multidrug resistant. The resistance percentages of bacteria were calculated as: percentage (%) = [(Number of resistant faecal coliforms to an antibiotic/ total number of tested faecal coliforms) × 100].

### Phenotypic identification of the production of Metallo-beta-lactamase (MBL), ESBL and AmpC enzymes

Bacteria resistant to imipenem and/or meropenem were subjected to the Imipenem-EDTA double-disk synergy test as described previously (47). Isolates resistant to cefotaxime and/or ceftazidime were subjected to ESBL testing following the CLSI guidelines and AmpC testing using phenylboronic acid (45, 48).

### Identification of antibiotic resistance genes and bacterial species

Putative MBL producing carbapenem resistant isolates (identified using the imipenem-EDTA double-disk synergy test) were subjected to multiplex PCR to identify the carbapenem resistance genes. The primer sets included bla_GES_, bla_GIM_, bla_IMI_, bla_IMP_, bla_KPC_, bla_NDM_, bla_OXA-23_, bla_OXA-40_, bla_OXA-48_, bla_OXA-51_, bla_OXA-58_, bla_VIM_ (Table 1) (49). Isolates displaying a positive ESBL phenotype and were phenotypically negative for AmpC production were further analysed using the ESBL multiplex-PCR. The primer sets were used to detect the *bla*_TEM_, *bla*_SHV_, and *bla*_CTX-M-_group genes 1, 2, 8, 9, and 25 (Table 1) (50, 51).

**Table 1:**
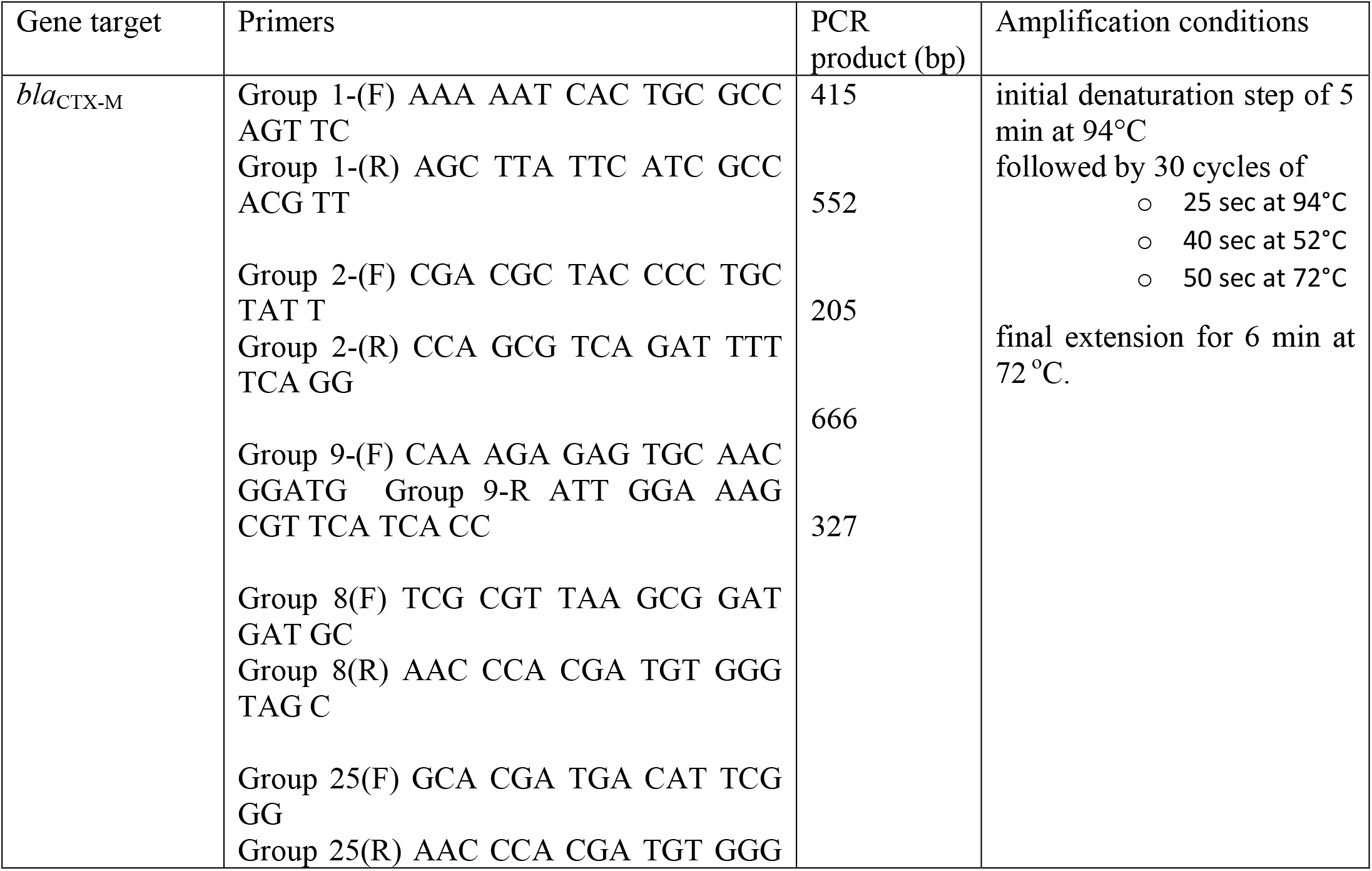
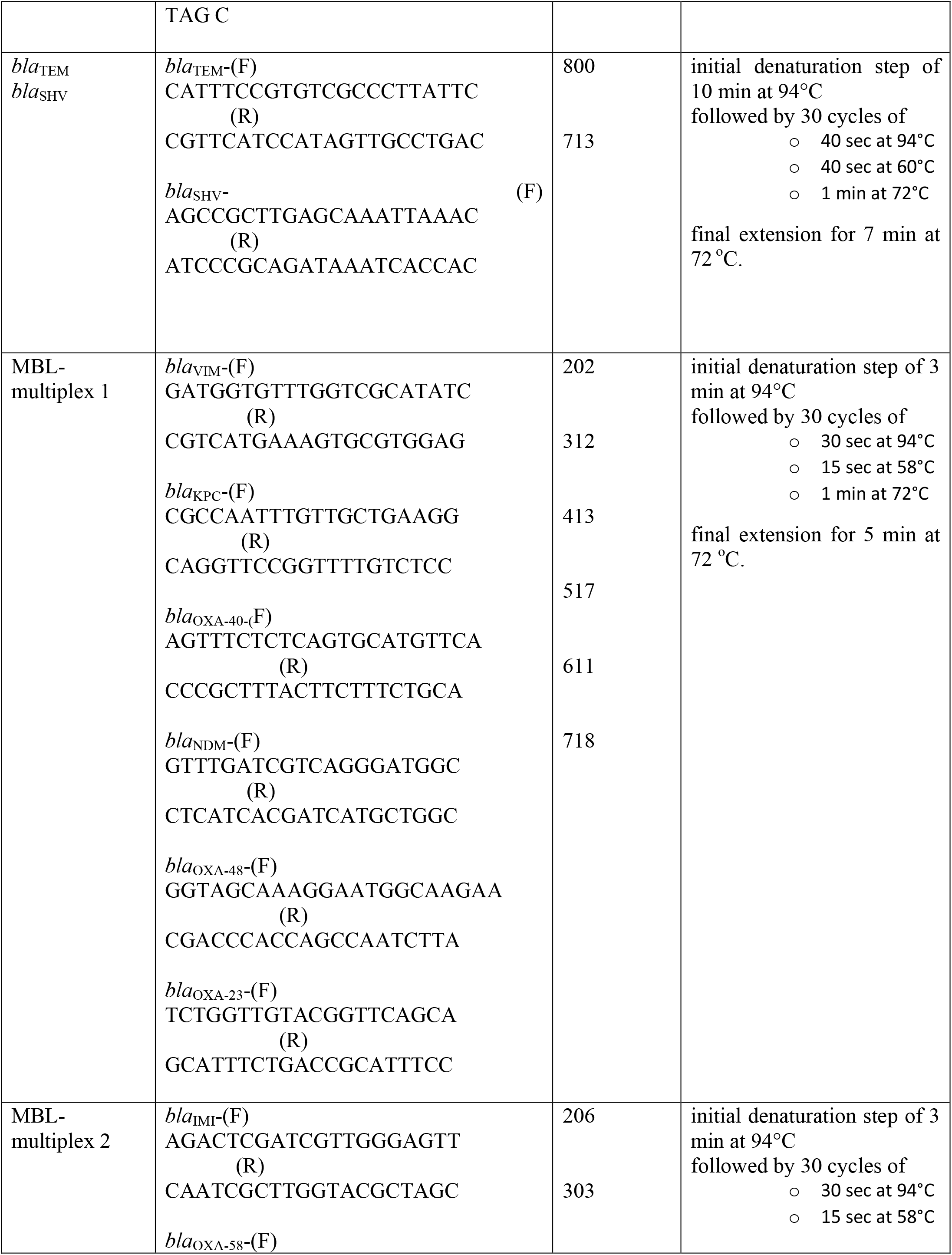
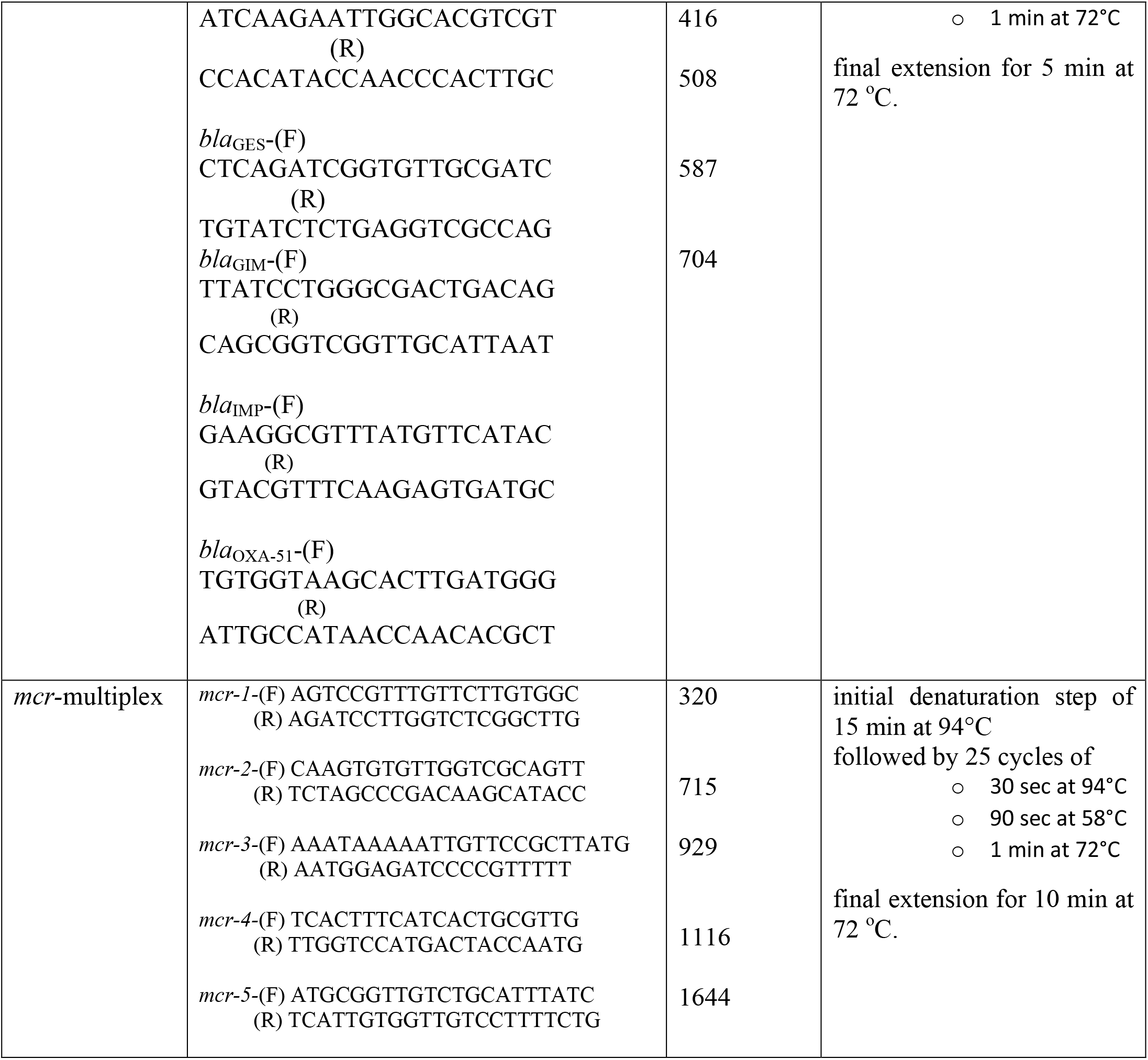
Primers used in multiplex PCRs to detect ESBL (50, 51), MBL (49) and *mcr*-genes (52)

Bacterial isolates resistant to colistin were further analysed by for the presence of the plasmid mediated colistin resistance mechanisms. Five primer sets were used to screen for the presence of *mcr*-1, 2, 3, 4 and 5, as recommended by the EU reference laboratory-antimicrobial resistance (Table 1) (52). The bacterial species of isolates carrying ESBL genes was identified by Sanger sequencing of the V3 and V4 region of bacterial 16S rRNA using forward primer: 5’-TCGTCGGCAGCGTCAGATGTGTATAAGAGACAGCCTACGGGNGGCWGCAG-3’ and reverse primer: 5’-GTCTCGTGGGCTCGGAGATGTGTATAAGAGACAGGACTACHVGGGTATCTAATC C-3’ (53).

### Plasmid transfer by transformation and replicon typing

Plasmids were extracted from ESBL positive isolates carrying the *bla*_TEM_, *bla*_SHV_, and *bla*_CTX-M_ genes using the Macherey-Nagel NucleoSpin plasmid isolation kit. The plasmids were transformed into *Escherichia coli* DH5α using heat-shock transformation (54, 55). The transformants were selected on LB agar supplemented with ampicillin (32 mg/L). The presence of *bla*_TEM_, *bla*_SHV_, and *bla*_CTX-M_ genes in transformants were confirmed by PCR. All transformants were subjected to antimicrobial susceptibility testing against imipenem, meropenem, ertapenem, ciprofloxacin, chloramphenicol, tetracycline, amikacin, gentamycin, kanamycin, trimethoprim and colistin. Replicon typing via PCR were performed on ESBL transformants with 18 pairs of primers recognizing FIA, FIB, FIC, HI1, HI2, I1-Iγ, L/M, N, P, W, T, A/C, K, B/O, X, Y, F, and FIIA in 3 multiplex panels (56).

## Results

### Antibiotic susceptibility patterns

In total, 498 faecal coliforms were isolated from all WWTP effluent samples, comprising 226 isolates from WWTP A and 272 from WWTP B. These isolates were subjected to antibiotic susceptibility testing. Among the tested β-lactam antibiotics, more than 90% of bacteria isolated from the two WWTPs were resistant to amoxicillin and ampicillin (Table 2, Figure 1); greater than 20% were resistant to cefotaxime or ceftazidime. All ceftazidime resistant isolates were also resistant to cefotaxime. Carbapenem resistance was detected at relatively lower levels (Table 2). Colistin resistance was found at a higher percentage in WWTP B effluent than in WWTP A. We also identified that there were no differences in the identification of colistin resistant isolates in antibiotic susceptibility testing by the agar dilution method compared to the microbroth dilution method. Multi-drug resistant (MDR) faecal coliforms were detected at approximately 50 % of the total isolates tested (Table 2). The resistance prevalence to other antibiotics were found at similar levels between the two WWTPs.

**Figure 1:**
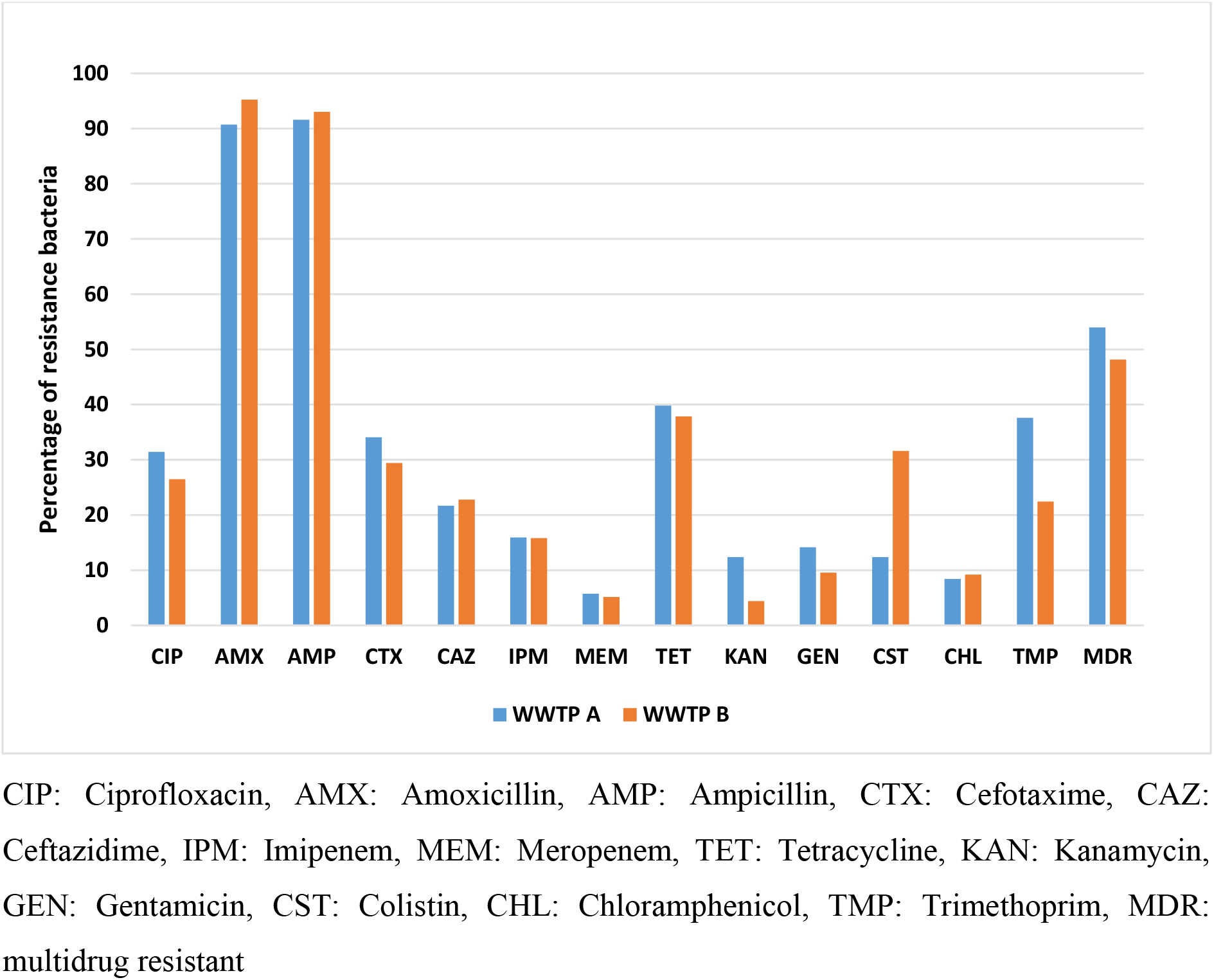
Percentage of faecal coliforms isolated from WWTP effluent samples identified with antibiotic resistance phenotypes.

**Table 2:** Prevalence of antibiotic resistance phenotypes in faecal coliforms isolated from WWTP effluent samples

| Antibiotic | Antibiotic-resistant faecal coliforms in WWTP effluent samples |  |  |  |
| --- | --- | --- | --- | --- |
|  | WWTP effluent A<br>(Total number of isolates =226) |  | WWTP effluent B<br>(Total number of isolates =272) |  |
|  | Number of resistant isolates | Percentage resistant (%) | Number of resistant isolates | Percentage resistant (%) |
| **CIP** | 71 | 31.42 | 72 | 26.47 |
| **AMX** | 205 | 90.71 | 259 | 95.22 |
| **AMP** | 207 | 91.59 | 253 | 93.01 |
| **CTX** | 77 | 34.07 | 80 | 29.41 |
| **CAZ** | 49 | 21.68 | 62 | 22.79 |
| **IPM** | 36 | 15.93 | 43 | 15.81 |
| **MEM** | 13 | 5.75 | 14 | 5.15 |
| **TET** | 90 | 39.82 | 103 | 37.87 |
| **KAN** | 28 | 12.39 | 12 | 4.41 |
| **GEN** | 32 | 14.16 | 26 | 9.56 |
| **CST** | 28 | 12.39 | 86 | 31.62 |
| **CHL** | 19 | 8.41 | 25 | 9.19 |
| **TMP** | 85 | 37.61 | 61 | 22.43 |
| **MDR** | 122 | 53.98 | 131 | 48.16 |

CIP: Ciprofloxacin, AMX: Amoxicillin, AMP: Ampicillin, CTX: Cefotaxime, CAZ: Ceftazidime, IPM: Imipenem, MEM: Meropenem, TET: Tetracycline, KAN: Kanamycin, GEN: Gentamicin, CST: Colistin, CHL: Chloramphenicol, TMP: Trimethoprim, MDR: multidrug resistant

### Detection of antibiotic resistance genes and speciation

Faecal coliforms with resistance to cefotaxime and ceftazidime were considered putative ESBL-producers (n = 157). AmpC production was confirmed in 26 of these isolates (13 from WWTP A and 13 from WWTP B). Of the 131 remaining isolates, 89 (39 from WWTP A and 50 from WWTP B) were identified as phenotypic ESBL producers. The ESBL multiplex PCR revealed the presence of *bla*_TEM_, *bla*_SHV-12_ and *bla*_CTX-M_ group 1 genes in 62 isolates. Among them, 49 isolates carried *bla*_TEM_ (25 from WWTP A and 24 from WWTP B), 7 *bla*_SHV-12_ (3 from WWTP A and 4 from WWTP B), 1 *bla*_CTX-M-1_ (WWTP B) and 5 *bla*_CTX-M-15_ (2 from WWTP A and 3 from WWTP B) (Table 3). Almost all ESBL producers were *E. coli* (46 carrying *bla*_TEM_, 2 carrying *bla*_SHV-12,_ 1 carrying *bla*_CTX-M-1_ and 2 carrying *bla*_CTX-M-15_). In addition, 7 isolates were *Klebsiella spp*. (5 carrying *bla*_SHV_, 2 carrying *bla*_CTX-M-15_) and 1 *bla*_CTX-M-15_ positive *Enterobacter spp*.

**Table 3:** Extended-spectrum β- lactamase genes identified in faecal coliforms isolated from WWTP effluent samples

| **ESBL gene** | **WWTP A**<br>**Number of ESBL**<br>**isolates** | **WWTP B**<br>**Number of ESBL**<br>**isolates** |
| --- | --- | --- |
| *bla*<sub>TEM-116</sub> | 6 | 5 |
| *bla*<sub>TEM-12</sub> | 3 | 2 |
| *bla*<sub>TEM-1-like</sub> (>99% identical to *bla*<sub>TEM-143</sub> or *bla*<sub>TEM-164</sub>) | 16 | 17 |
| *bla*<sub>CTX-M-1</sub> | 0 | 1 |
| *bla*<sub>CTX-M-15</sub> | 2 | 3 |
| *bla*<sub>SHV-12</sub> | 3 | 4 |

In total, 79 isolates (36 from WWTP A and 43 from WWTP B) resistant to imipenem were screened for MBL production. Metallo-beta-lactamase production was identified in 36 isolates. However, all isolates were negative for known MBL genes, using the MBL multiplex PCR. More than 100 colistin resistant isolates were detected. However, none of these isolates were positive for the *mcr*-genes using the *mcr*-targeted PCR analysis.

### Plasmid transformation, plasmid resistance profile and replicon typing

The plasmids were extracted from 62 ESBL faecal coliform isolates (30 from WWTP A and 32 from WWTP B). Plasmids extracted from 52 ESBL isolates (46 plasmids carried *bla*_TEM_, 2 plasmids *bla*_CTX-M15_, 1 *bla*_CTX-M-1_ and 3 with *bla*_SHV-12_) were successfully transferred into *E. coli* Dh5α recipients. The plasmids extracted from the 10 ESBL isolates (3 with *bla*_TEM_, 3 with *bla*_CTX-M15_ and 4 with *bla*_SHV-12_) could not be transferred, suggesting that the ESBL genes identified in these isolates are located on the bacterial chromosomes.

All transformants were sensitive to the tested carbapenem antibiotics (imipenem, meropenem and ertapenem) and gentamicin, 16 were resistant to chloramphenicol, 16 resistant to tetracycline, 10 resistant to trimethoprim, 3 resistant to colistin, 2 resistant to kanamycin, 1 resistant to amikacin and 1 resistant to ciprofloxacin. Nine transformants show a resistance phenotype to two drugs from different classes and eight show a multidrug resistance phenotype. The plasmids from 21 transformants could be typed using PCR replicon typing. Of these, 11 plasmids were IncF replicon type (3 IncFIA, 1 IncFIIA, 1 IncFIB and 6 non-specific IncF group), 5 were IncI1 replicon type, 2 were IncHI1 replicon type, 2 were IncHI2 replicon type and 1 was IncA/C replicon type. The IncF group of replicons was the most prevalent across plasmids from both WWTPs. The IncI1 type was detected only in bacteria isolated from the WWTP B effluent samples.

## Discussion

Our work presents the antibiotic resistance patterns of faecal coliforms in the effluent of two WWTPs in Ireland. Wastewaters with faecal contamination are considered reservoirs for ARB and ARGs in the environment (57, 58). Among all tested antibiotics, resistance to amoxicillin and ampicillin was most prevalent, followed by tetracycline and, cefotaxime and ciprofloxacin. High levels of β-lactam resistance were previously detected in Enterobacteriaceae in an urban WWTP (59), and resistance to tetracycline and fluoroquinolones was found at lower rates. The higher resistance rate of *E. coli* to ampicillin and tetracycline as well as lower rates of resistance to ciprofloxacin were detected in WWTPs in Portugal (60). A study of raw sewage in Brazil identified 100% sensitivity of *E. coli* to ciprofloxacin and amoxicillin and tetracycline resistance levels in the range of 50 – 75% (61). The proportions of ciprofloxacin resistant faecal coliforms were 31.42% in WWTP A effluent and 26.47% in WWTP B. This resistance rate is higher in comparison to reported levels of ciprofloxacin resistance in the *E. coli* isolated from other WWTPs (59, 62, 63). In general, differences in antibiotic resistance percentages were observed between the two WWTPs, particularly for colistin, trimethoprim and kanamycin (Figure 1). As these two WWTPs are using the same treatment process, this difference may be associated with their location or other external factors.

The presence of colistin resistant and carbapenem resistant isolates in tested WWTP effluents raises the possibility of transferability or risk to human health. These antibiotics are known as ‘last resort’ antibiotics to treat MDR bacteria. Previous studies on colistin resistant Enterobacteriaceae focus mainly on food, human and animal samples (64–67). To date, there are only a few studies conducted on the identification of *mcr* genes in waterborne bacteria (68–72). 114 faecal coliform isolates resistant to colistin (28 from WWTP A and 86 from WWTP B) were detected in our work, none of them were positive for the *mcr* genes. Among colistin resistant isolates, the plasmids extracted from 43 isolates (4 from WWTP A and 39 from WWTP B) were successfully transferred to the recipient *E. coli* DH5α in the transformation studies. The proportion of colistin resistant coliforms in our study was lower than those in the studies of Igwaran et al.(68), where they found approximately 60% of isolates with resistance to colistin. Carbapenem resistant Enterobacteriaceae were studied in hospital wastewater and in WWTPs previously (73–76). The percentage of carbapenem resistance phenotypes in our work were considered high in comparison to previous studies in wastewater (33, 77). However, there were no known carbapenem resistance genes detected in the carbapenem resistant coliforms. This suggests that other resistance mechanisms are responsible for the resistance phenotypes and thus require further study to characterise these novel mechanisms.

Multi-drug resistant faecal coliforms were retrieved at high rates from all effluent samples. In the study of Lefkowitz and Durán (78), 60% of *E. coli* in WWTP effluent were resistant to two or more antibiotics and 25% to four or more antibiotics. The study of Garcia *et al* (79) in WWTP effluent showed no more than 12% of *E. coli* were resistant to two antibiotics and less than 10% to three or more antibiotics. *Escherichia coli* (34.3%) were found to be resistant to two or more antibiotics and 8.8% to four or more antibiotics in treated wastewater in Portugal (59). The MDR faecal coliforms in our study were found in the same range of the study of Lefkowitz and Durán, but at a higher percentage than in others.

The ESBLor AmpC producing faecal coliforms were recovered from all WWTP effluent samples. The rate of ESBL producing *Enterobacteriacea* in our work were within the range of previous studies. It was considerably high in comparison to some studies of WWTP effluent (0.5-9.8%) (34, 80-83). However it was lower than those studies in wastewater (45-100%) (84, 85). The high load of bacteria and rich nutrient environment in WWTPs facilitates the transfer of ARGs among bacteria (11, 12). These may explain the relatively high rates of ESBL producers in WWTP effluent.

The *bla*_TEM_ were the most prevalent beta-lactamase in this study, which is similar to a study on hospital wastewater in Brazil (85). However, this is in contrast with other findings with *bla*_CTX-M_ being the most frequent ESBL genes from various samples including in hospital effluent, surface water and WWTPs (34, 39, 86). This result was confirmed by qPCR data performed in another study in the framework of the StARE project which showed the detection of *bla*_TEM_ genes, where *bla*_SHV_ and *bla*_CTX-M_ group 1 were not detected (22). Most of ESBL genes were found in *E. coli*, others were carried by *Klebsiella spp.* in our work. These results are in an agreement with previous findings which indicated that *E. coli* were the most common ESBL-producers among Enterobacteriaceae (39).

Transformation of plasmids carrying ESBL genes were successful for 88% of ESBL donor isolates in this work. Different replicon variants were found in the ESBL plasmids. The most prevalent replicon was IncF group which were also reported in other studies of plasmid replicon typing in Enterobacteriaceae (87–89). These replicons have a narrow host range and can be transfer easily among *E. coli* species (90). The cross-resistance of ESBL producers to other antibiotic causes of great concern as ESBL genes are frequently located on conjugative plasmids carrying other ARGs (91). In this work eight of 52 transferable plasmids carrying ESBL genes expressed a MDR phenotype.

## Conclusion

Effluent samples from two WWTPs demonstrated the presence of ARB and MDR and, of particular importance, a source of a relatively high proportion of ESBL-producing, carbapenem and colistin resistant Enterobacteriaceae. Although the bacteria were phenotypically resistant to colistin or carbapenems no known mobile resistance mechanisms were detected, despite the ability to transfer the resistance phenotype via transformation. Thus, faecal coliforms from WWTP effluent are sources of novel ARGs conferring resistance to antibiotics of last line of defence. The ability of these bacteria to survive in water has been demonstrated for many years. The significance of this study is the identification of the role of WWTPs as a potential control point to reduce or stop the movement of resistant bacteria and genes into the environment from further upstream sources, such as human or animal waste. In addition, this enables the use of additional treatment technologies to be added to WWTPs to stop or reduce such ARB and ARGs entering the water environments globally.

## Acknowledgments

This work was funded by the Irish Environmental Protection Agency, in the frame of the collaborative international consortium of Water challenges for a changing world Joint Programming Initiative (Water JPI) Pilot Call, project Stare and the EPA – Maynooth University co-fund project “Survival of mobile antibiotic resistance in water”. COST Action ES1403: New and emerging challenges and opportunities in wastewater reuse (NEREUS).

